# The scaffold-dependent function of RIPK1 in dendritic cells promotes injury-induced colitis

**DOI:** 10.1101/2021.02.27.433202

**Authors:** Kenta Moriwaki, Christa Park, Kazuha Koyama, Sakthi Balaji, Kohei Kita, Ryoko Yagi, Sachiko Komazawa-Sakon, Tatsuya Asuka, Hiroyasu Nakano, Yoshihiro Kamada, Eiji Miyoshi, Francis K.M. Chan

## Abstract

Receptor interacting protein kinase 1 (RIPK1) is a cytosolic multidomain protein that controls cell life and death. While RIPK1 promotes cell death through its kinase activity, it also functions as a scaffold protein to promote cell survival by inhibiting FADD-caspase 8-dependent apoptosis and RIPK3-MLKL-dependent necroptosis. This pro-survival function is highlighted by excess cell death and a perinatal lethality in *Ripk1*^−/−^ mice. Recently, loss of function mutation of *RIPK1* was found in patients with immunodeficiency and inflammatory bowel diseases. Hematopoietic stem cell transplantation restored not only immunodeficiency but also intestinal inflammatory pathology, indicating that RIPK1 in hematopoietic cells is critical to maintain intestinal immune homeostasis. Here, we generated dendritic cell (DC)-specific *Ripk1*^−/−^ mice in a genetic background with loss of RIPK1 kinase activity and found that the mice developed spontaneous colonic inflammation characterized by increased neutrophil infiltration. In addition, these mice were highly resistant to injury-induced colitis. The increased neutrophil infiltration in the colon and the resistance to colitis were restored by dual inactivation of RIPK3 and FADD, but not by inhibition of RIPK3, MLKL, or ZBP1 alone. Altogether, these results reveal a scaffold activity-dependent role of RIPK1 in protecting colonic DCs from apoptotic insults and maintenance of colonic immune homeostasis.

## Introduction

Receptor interacting protein kinase 1 (RIPK1) is a cytosolic serine/threonine kinase that functions downstream of various cell surface immune-related receptors such as tumor necrosis factor receptor 1 (TNFR1)^1^. RIPK1 consists of a kinase domain (KD) and a death domain (DD) at N- and C-termini respectively. In addition, RIPK1 harbors the RIP homotypic interaction motif (RHIM) in the intermediate region. Upon tumor necrosis factor (TNF) stimulation, RIPK1 is recruited to TNFR1 through homotypic DD interaction. In the membrane-bound TNFR1 complex, called complex I, RIPK1 is modified by K63- and M1-ubiquitination by the action of cellular inhibitor of apoptosis 1 (cIAP1) and linear ubiquitin chain assembly complex (LUBAC)^2^. These ubiquitin chains on RIPK1 act as a scaffold to recruit various signaling kinases such as an inhibitor of κB kinase (IKK) α/β, IKKε, transforming growth factor β activated kinase 1 (TAK1), and TANK binding kinase 1 (TBK1)^3^. These kinases directly or indirectly phosphorylate RIPK1 to inhibit its activation. In addition, IKKs phosphorylate inhibitor of κB (IκB), leading to NF-κB-dependent expression of pro-survival molecules such as cellular FLICE inhibitory protein (cFLIP)^4^. When these kinases are inhibited, RIPK1 interacts with FADD and caspase 8 to induce RIPK1-dependent apoptosis. In addition, when expression of pro-survival proteins is inhibited by, for example, cycloheximide, cells can undergo RIPK1-independent apoptosis. This RIPK1-indepenent apoptosis is potentiated by loss of RIPK1 because of less induction of pro-survival gene expression.

Besides apoptosis, RIPK1 also plays a role in an alternative cell death modality termed necroptosis, a regulated form of cell necrosis mediated by RIPK3 and the downstream effector mixed lineage kinase domain-like pseudokinase (MLKL)^5^. During TNFR1-induced necroptosis, RIPK1 interacts with RIPK3 via homotypic RHIM interaction in a RIPK1 kinase activity-dependent manner. The interaction promotes activation of RIPK3 and the conversion of the RIPK1-RIPK3 complex into an amyloid signaling complex called the necrosome ^6, 7^. In addition to RIPK1, RIPK3 can also be activated by interaction with other RHIM-containing proteins such as the toll like receptor adaptor protein TRIF and the cytosolic Z-RNA receptor ZBP1^8, 9^. Paradoxically, RIPK1 has been discovered to inhibit TRIF-RIPK3 and ZBP1-RIPK3 necrosomes in a RHIM-dependent and kinase activity-independent manner under certain conditions ^10–13^. Therefore, the scaffold and the kinase functions of RIPK1 promotes cell survival and cell death, respectively.

It has been demonstrated by various genetic and chemical approaches that RIPK1 kinase activity is involved in the development of infectious and non-infectious diseases^14^. Since *Ripk1* kinase-dead mutant knock-in mice develop and grow normally without any obvious abnormality, RIPK1 has garnered attention as a therapeutic target for various inflammatory diseases^15^. In contrast, *Ripk1*^−/−^ mice develop systemic inflammation and an early postnatal lethality due to aberrant apoptosis and necroptosis in multiple tissues^10, 11, 16^. These phenotypes are rescued by concomitantly blocking apoptosis through deletion of either FADD or caspase 8 and necroptosis through deletion of either RIPK3 or MLKL. Recently, biallelic loss of function mutations of *RIPK1* were found in patients suffered from primary immunodeficiency and peripheral inflammation^17–19^. One of the most common pathologies observed in the patients was early-onset inflammatory bowel disease. Interestingly, hematopoietic stem cell transplantation resolved clinical symptoms of inflammatory bowel disease as well as reduced a frequency of recurrent infection, highlighting that RIPK1 in hematopoietic cells is critical to maintain immune homeostasis and protect autoinflammation in peripheral tissues^17^. Although cell type-specific functions of RIPK1 have been investigated using conditional *Ripk1* knockout mice^20–24^, the immune cell type in which RIPK1 functions to protect against inflammatory bowel disease is unknown at present.

Dendritic cells (DCs) are central immune effectors that recognize foreign antigens and control acquired and innate immune responses^25^. We previously reported that RIPK3 promotes cytokine production and tissue repair in a kinase activity-independent manner in DCs during injury-induced colitis ^26–28^. By contrast, it remains unknown whether RIPK1 in DCs plays a similar role in intestinal inflammation. Here, we utilized *Ripk1* kinase-dead floxed mice and investigated the differential roles of RIPK1 kinase and scaffold functions in DCs in intestinal immune homeostasis. We found that DC-specific deletion of RIPK1 caused FADD-dependent spontaneous colonic inflammation characterized by increased neutrophil infiltration. Despite the elevated basal colonic inflammation, loss of RIPK1 in DCs surprisingly rendered mice resistant to injury-induced colitis. These results demonstrate that RIPK1 in DCs has a critical impact on intestinal immune homeostasis.

## Results

### DC-specific deletion of RIPK1 caused splenic inflammation independent of RIPK1 kinase activity

RIPK1 controls multiple signaling pathways through kinase activity-dependent and - independent functions. To delineate the kinase activity-dependent and -independent roles in DCs, we crossed *Ripk1* kinase-dead mutant knock-in mice (*Ripk1*^kd/kd^)^29^, in which exon 4 was flanked by loxP sequences, with CD11c-Cre transgenic mice (CD11c;*Ripk1*^kd/kd^ mice). In the spleen, DCs are mainly divided into two subsets: CD8α^+^CD11b^−^ conventional DC1 (cDC1) and CD8α^−^ CD11b^+^ cDC2^25^. RIPK1 expression was slightly higher in cDC1 than in cDC2 (Fig. 1a). In both subsets, RIPK1 was completely deleted in CD11c;*Ripk1*^kd/kd^ mice (Fig. 1b). Previous reports showed that *Ripk1*^kd/kd^ mice grow normally and did not show any overt abnormality at steady state^29, 30^. We confirmed normal spleen size and splenic immune cell population (Fig. 1c, d). In contrast, we observed splenomegaly and significant increase of neutrophil and macrophage numbers in the spleen of CD11c;*Ripk1*^kd/kd^ mice (Fig. 1e, f). Similarly, the increased numbers of these cells were observed in the bone marrow of CD11c;*Ripk1*^kd/kd^ mice (Fig. 1g). Although deletion of RIPK1 sensitizes cells to apoptosis and necroptosis, the number of splenic and bone marrow DCs did not alter in CD11c;*Ripk1*^kd/kd^ mice (Fig. 1f, g). This might be due to a compensatory supply from progenitors to maintain DC number.

**Fig. 1.**
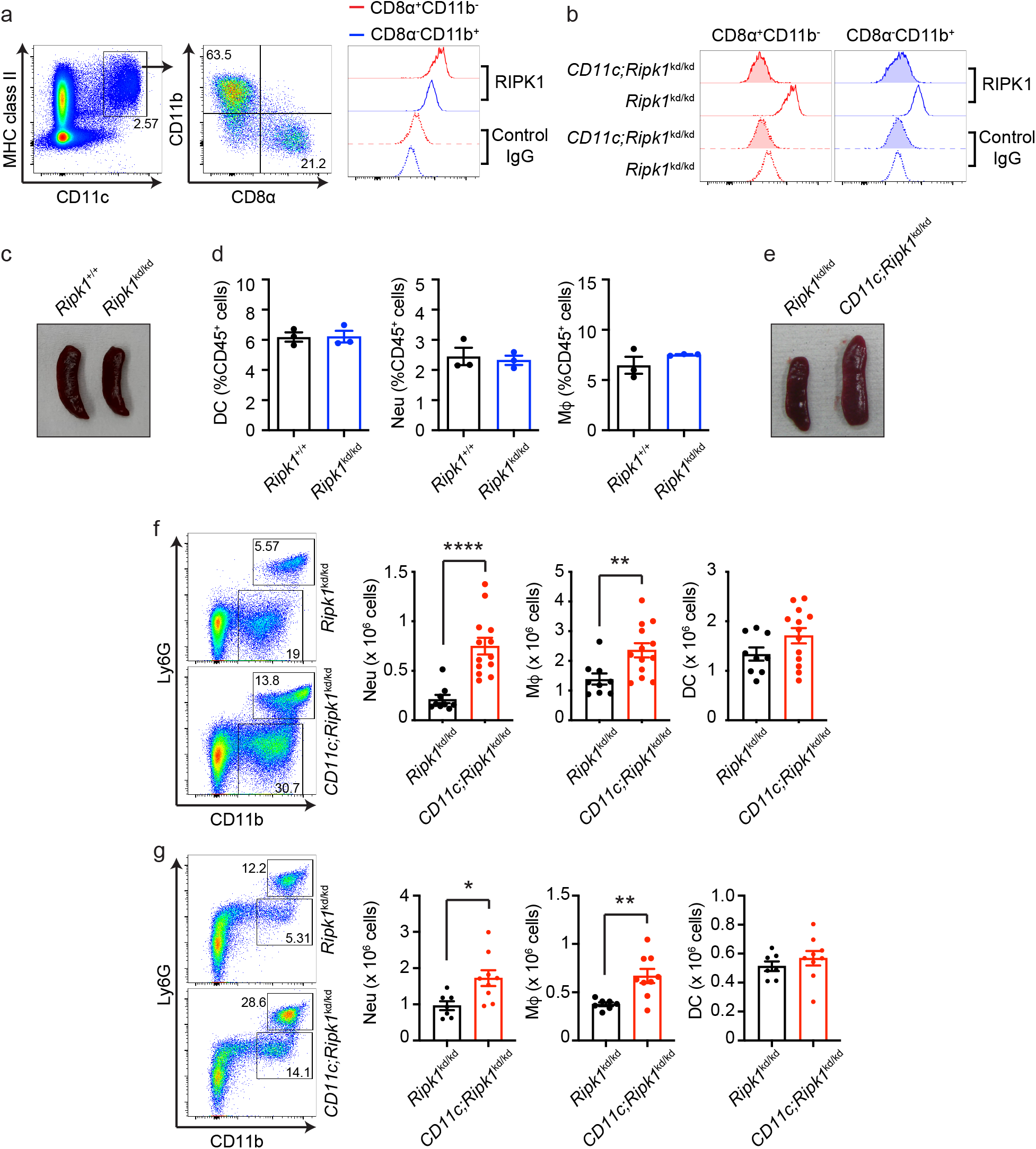
DC-specific RIPK1 deletion caused inflammation in the absence of RIPK1 kinase activity. **a,b** Splenocytes of *Ripk1*^kd/kd^ (**a, b**) and CD11c;*Ripk1*^kd/kd^ mice (**b**) were subjected to RIPK1 intracellular staining. Representative results of flow cytometry are shown (n = 4 mice). Live/Dead aqua^−^CD45^+^ cells are presented (**a**, the most left panel). **c, e** Representative pictures of the spleen from mice with the indicated genotypes are shown. **d** Percentages of CD3^−^CD19^−^ CD11c^+^ DCs, CD3^−^CD19^−^CD11c^−^CD11b^+^Ly6G^+^ neutrophils (Neu), and CD3^−^CD19^−^CD11c^−^ CD11b^+^Ly6G^−^ macrophages (Mφ) among CD45^+^ cells are shown. Results are mean ± SEM (n = 3 per each genotype). **f, g** The number of CD45^+^CD3^−^B220^−^CD11c^+^ DCs, CD45^+^CD3^−^B220^−^ CD11c^−^CD11b^+^Ly6G^+^ neutrophils (Neu), and CD45^+^CD3^−^B220^−^CD11c^−^CD11b^+^Ly6G^−^ macrophages (Mφ) in the spleen (**f**) and bone marrow (**g**) are shown. Results are mean ± SEM (**f:** n = 9 for *Ripk1*^kd/kd^ mice and n = 13 for CD11c;*Ripk1*^kd/kd^ mice, **g:** n = 7 for *Ripk1*^kd/kd^ mice and n = 9 for CD11c;*Ripk1*^kd/kd^ mice). Mice from the intercross between *Ripk1*^+/−^ mice were used in **c** and **d**. Mice from the intercross between *Ripk1*^kd/kd^ and CD11c;*Ripk1*^kd/kd^ mice were used in **a**, **b**, and **e-g**. \**p* < 0.05, \*\**p* < 0.01, \*\*\*\**p* < 0.0001 (unpaired t test with Welch’s correction).

### DC-specific deletion of RIPK1 caused spontaneous colonic inflammation

In the colon, DCs are divided into three groups: CD103^+^CD11b^−^, CD103^+^CD11b^+^, and CD103^−^ CD11b^+^ DCs. These cells coordinately control colonic immune homeostasis, allowing an optimal response to commensal bacteria and food antigens^31^. RIPK1 was equally expressed in these three subsets (Fig. 2a). While RIPK1 was deleted in CD103^+^CD11b^−^ and CD103^+^CD11b^+^ DCs in CD11c;*Ripk1*^kd/kd^ mice, it still expressed in CD103^−^CD11b^+^ DCs (Fig. 2b). This is most likely due to much lower CD11c expression in CD103^−^CD11b^+^ DCs than in the other two subsets (Fig. 2c). As in the spleen, we found increased neutrophil infiltration in the colon of CD11c;*Ripk1*^kd/kd^ mice, although microscopically visible tissue damage was not observed (Fig. 2d-f). The increased colonic neutrophil infiltration was also observed in CD11c;*Ripk1*^fl/fl^ mice (Fig. 2g). In contrast, there was no increase in neutrophils in the colon of *Ripk1*^kd/kd^ mice compared to *Ripk1*^+/+^ mice (Fig. 2h). In addition, neutrophil chemokines *Cxcl1* and *Cxcl2* were highly expressed in the colon of CD11c;*Ripk1*^kd/kd^ mice (Fig. 2i). Unlike in the spleen, the number of DCs and macrophages was reduced in the colon (Fig. 2j). Consistent with the efficiency of RIPK1 deletion (Fig. 2b), the number of CD103^+^CD11b^−^ and CD103^+^CD11b^+^ DCs were more strongly reduced compared with that of CD103^−^CD11b^+^ DCs (Fig. 2k). The number of DCs and macrophages in the colon was normal in *Ripk1*^kd/kd^ mice (Fig. 2l). These results indicate that deletion of RIPK1 in DCs, especially in CD103^+^ DCs, caused spontaneous inflammation in the colon.

**Fig. 2.**
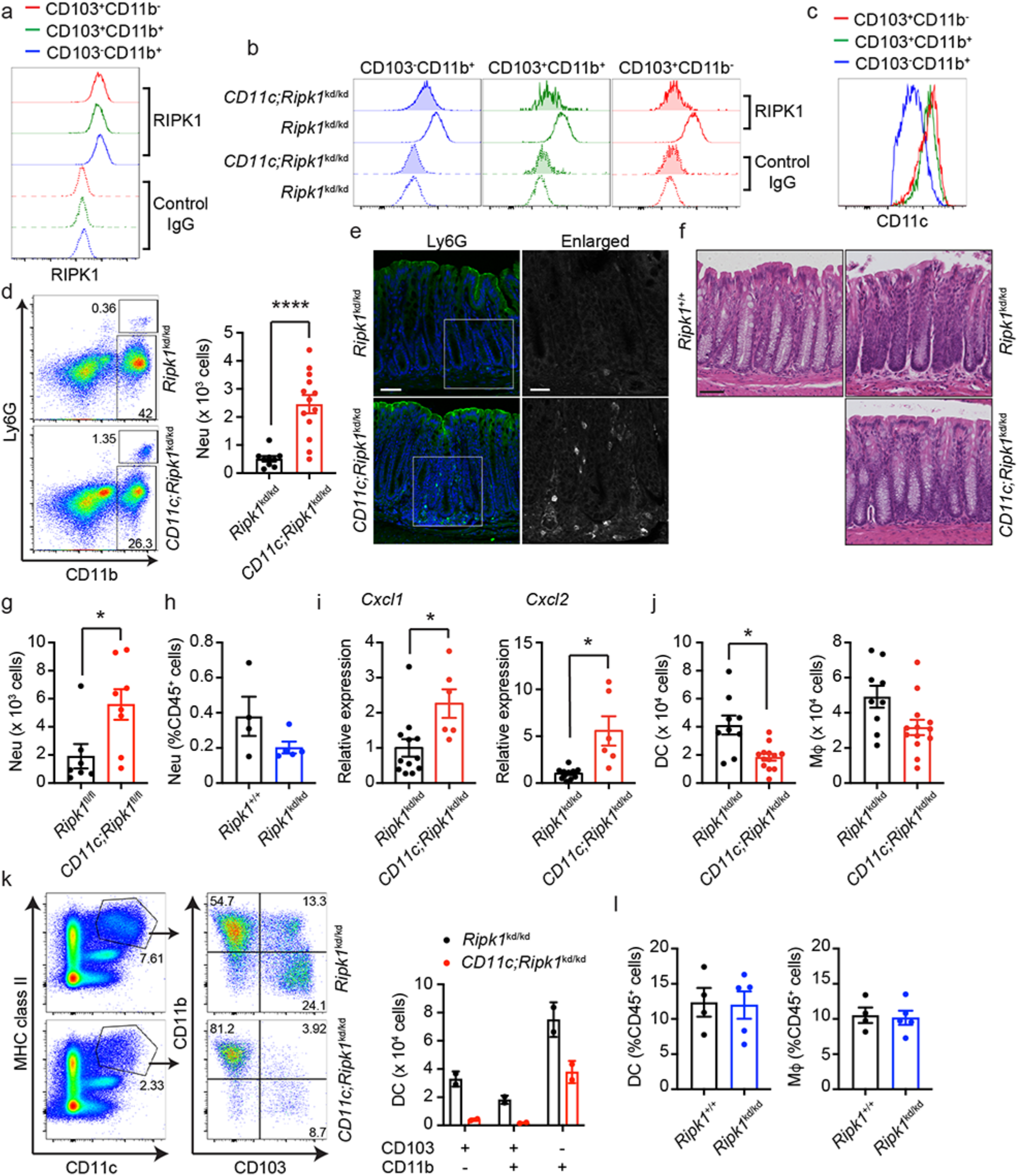
DC-specific RIPK1 deletion caused spontaneous neutrophil infiltration in the colon. **a,b** Colonic lamina propria cells isolated from of *Ripk1*^kd/kd^ (**a, b**) and CD11c;*Ripk1*^kd/kd^ mice (**b**) were subjected to RIPK1 intracellular staining. Representative results of flow cytometry are shown (n = 4 per each genotype). Live/Dead aqua^−^CD45^+^CD11c^+^MHC class II^+^ cells are presented. **c** Representative result of CD11c expression in three colonic DC subsets is shown (n = 4 per each genotype). **d, j** The number of colonic CD45^+^CD3^−^B220^−^CD11c^−^CD11b^+^Ly6G^+^ neutrophils (Neu) (**d**), CD45^+^CD3^−^B220^−^CD11c^+^ DCs, and CD45^+^CD3^−^B220^−^CD11c^−^ CD11b^+^Ly6G^−^ macrophages (Mφ) (**j**) are shown. Results are mean ± SEM (n = 9 for *Ripk1*^kd/kd^ mice and n = 13 for CD11c;*Ripk1*^kd/kd^ mice). **e** The colon was stained for Ly6G (green) (n = 2 per each genotype). The areas indicated by the white squares are enlarged and shown on the right. Nuclei (blue) were stained with DAPI. Scale bars: 50 μm (left panel) and 25 μm (right panel). **f** H& staining of the colon from mice with the indicated genotypes (n = 3 per each genotype). Scale bars: 50 μm. **g** The number of colonic CD45^+^CD3^−^CD19^−^CD11c^−^CD11b^+^Ly6G^+^ neutrophils (Neu) are shown. Results are mean ± SEM (n = 7 for *Ripk1*^fl/fl^ mice and n = 8 for CD11c;*Ripk1*^fl/fl^ mice). **h, l** Percentages of CD3^−^CD19^−^CD11c^−^CD11b^+^Ly6G^+^ neutrophils (Neu) (**h**), CD3^−^CD19^−^CD11c^+^ DCs, and CD3^−^CD19^−^CD11c^−^CD11b^+^Ly6G^−^ macrophages (Mφ) (**l**) are shown. All populations were gated on CD45+ cells. Results are mean ± SEM (n = 4 for *Ripk1*^+/+^ mice and n = 5 for *Ripk1*^kd/kd^ mice). **i** The expression of *Cxcl1* and *Cxcl2* in the colon was determined by real-time qPCR. Results are mean ± SEM (n = 12 for *Ripk1*^kd/kd^ mice and n = 6 for CD11c;*Ripk1*^kd/kd^ mice). **k** Colonic lamina propria cells were analyzed by flow cytometry. Representative results are shown in the left panels (n = 4 mice). Live/Dead aqua^−^CD45^+^ cells are presented. The number of three colonic DC subsets are shown in the graph on the right. Results are mean ± SEM (n = 2 per each genotype. Representative result from two independent experiments is shown.). Mice from intercross between *Ripk1*^fl/fl^ and CD11c;*Ripk1*^fl/fl^ mice were used in **g**. *Ripk1*^+/+^ (**f, h, l**) and *Ripk1*^kd/kd^ (**h, l**) mice were obtained from intercross between *Ripk1*^+/−^ mice. Mice from the intercross between *Ripk1*^kd/kd^ and CD11c;*Ripk1*^kd/kd^ mice were used in others. \**p* < 0.05, \*\*\*\**p* < 0.0001 (unpaired t test with Welch’s correction).

### Increased neutrophil in the colon of CD11c;Ripk1^kd/kd^ mice was dependent on FADD but not RIPK3

Deletion of RIPK1 sensitizes cells to FADD/Caspase 8-dependent apoptosis and RIPK3/MLKL-dependent necroptosis. To examine whether these types of cell death contribute to the spontaneous inflammation, we crossed CD11c;*Ripk1*^kd/kd^ mice with RIPK3 RHIM-deleted mice (*Ripk3*^ΔR/ΔR^ mice)^28^ and/or *Fadd*^−/−^ mice. *Ripk3*^ΔR/ΔR^ mice are resistant to necroptosis because the RHIM is an essential motif for necroptosis induction ^28^. Although *Fadd*^−/−^ mice are embryonic lethal due to excessive necroptosis^32^, they are viable in the *Ripk3*^ΔR/ΔR^ genetic background^28^. In the spleen, the increased infiltration of neutrophils and macrophages was almost completely restored by deletion of RIPK3 RHIM alone (Fig. 3a, 2^nd^ vs 4^th^ bars). In contrast, the number of neutrophils and DCs in the colon was restored by deletion of both FADD and RIPK3 RHIM but not either RIPK3 RHIM or MLKL alone (Fig. 3b, c). Similarly, the increased expression of neutrophil chemokines *Cxcl1* and *Cxcl2* were restored only when FADD and RIPK3 RHIM were deleted (Fig. 3d). These results suggest that RIPK1 deletion caused spontaneous inflammation by inducing necroptosis in the spleen and apoptosis in the colon.

**Fig.3.**
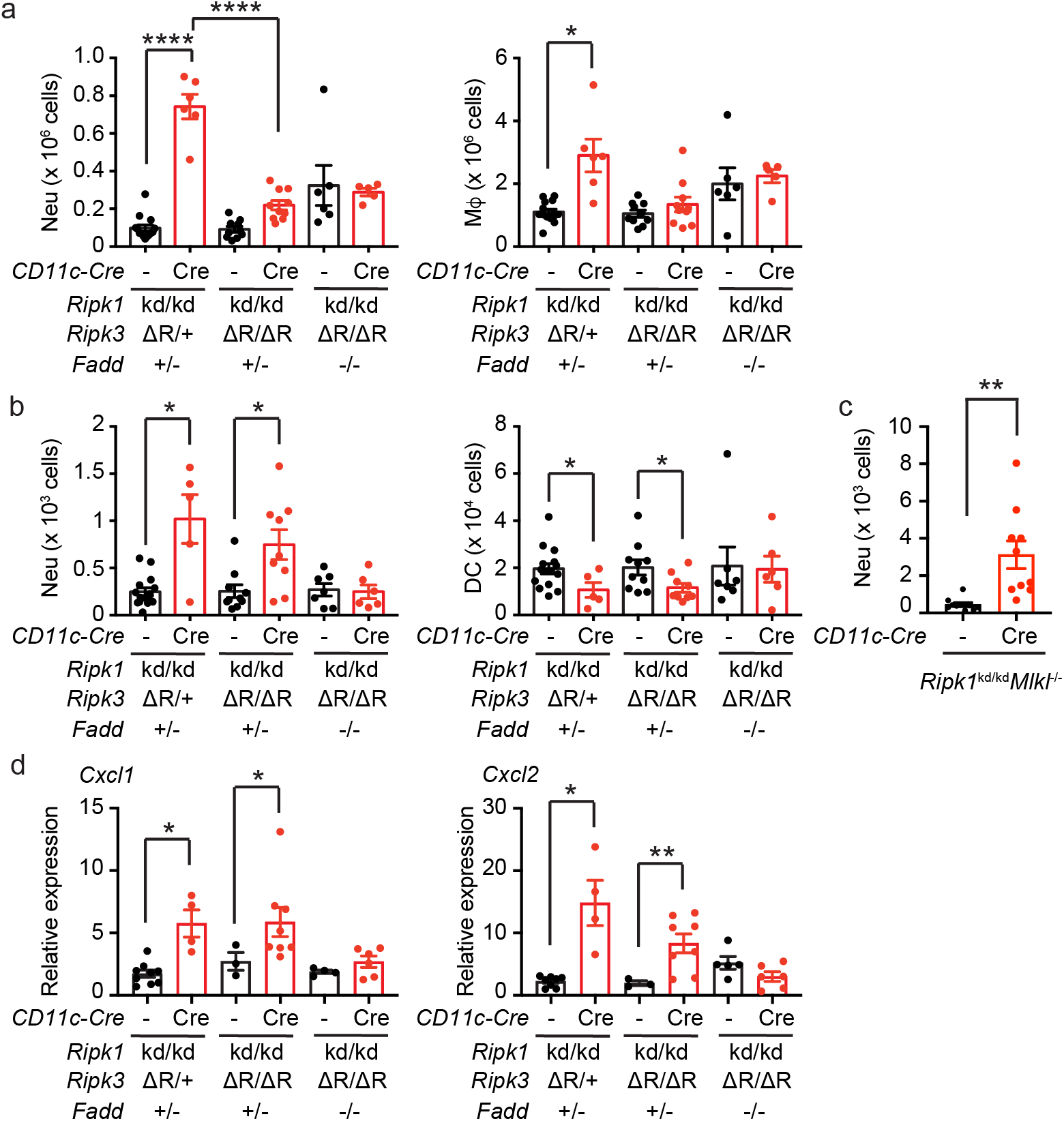
The increased neutrophil infiltration in the colon was restored by deletion of RIPK3 RHIM and FADD. **a-c** Splenocytes (**a**) and colonic lamina propria cells (**b, c**) isolated from mice with the indicated genotypes were analyzed by flow cytometry. The number of CD3^−^B220^−^ CD11c^−^CD11b^+^Ly6G^+^ neutrophils (Neu, **a** and **b**), CD3^−^CD19^−^CD11c^−^CD11b^+^Ly6G^+^ neutrophils (**c**), CD3^−^B220^−^CD11c^+^ DCs, and CD3^−^B220^−^CD11c^−^CD11b^+^Ly6G^−^ macrophages (Mφ) are shown. Results are mean ± SEM (n = 5-15 per each genotype). **d** The expression of *Cxcl1* and *Cxcl2* in the colon was determined by real-time qPCR. Results are mean ± SEM (n = 3-9 per each genotype). Mice from intercross between *Ripk1*^kd/kd^*Ripk3*^ΔR/ΔR^*Fadd*^−/−^ and CD11c;*Ripk1*^kd/kd^*Ripk3*^ΔR/+^*Fadd*^+/−^ mice were used in **a**, **b**, and **d**. Mice from intercross between *Ripk1*^kd/kd^*Mlkl*^−/−^ and CD11c;*Ripk1*^kd/kd^*Mlkl*^−/−^ mice were used in **c**. \**p* < 0.05, \*\**p* < 0.01, \*\*\*\**p* < 0.0001 (one-way ANOVA in **a** and unpaired t test with Welch’s correction in others).

### ZBP1 contributes to the spontaneous inflammation in the spleen, but not the colon

ZBP1 is a RHIM-containing protein that induces necroptosis through RHIM-dependent interaction with RIPK3 upon recognition of endogenous or viral Z-RNA^33–35^. RIPK1 deletion triggers ZBP1 and RIPK3-dependent necroptosis. As such, RIPK1 has a critical role in blocking ZBP1-RIPK3 interaction and necroptosis ^12, 13^. In the absence of ZBP1, the number of splenic neutrophils in CD11c;*Ripk1*^kd/kd^ mice was similar compared to *Ripk1*^kd/kd^ mice (Fig. 4a). In contrast, increased neutrophil infiltration in the colon of CD11c;*Ripk1*^kd/kd^ mice was not reversed by *Zbp1* deletion, although the difference in cell number was not statistically different between CD11c;*Ripk1*^kd/kd^;*Zbp1*^−/−^ and *Ripk1*^kd/kd^;*Zbp1*^−/−^ mice (Fig. 4b). These results further support the notion that necroptosis contributes to the spontaneous inflammation in the spleen, but not the colon.

**Fig.4.**
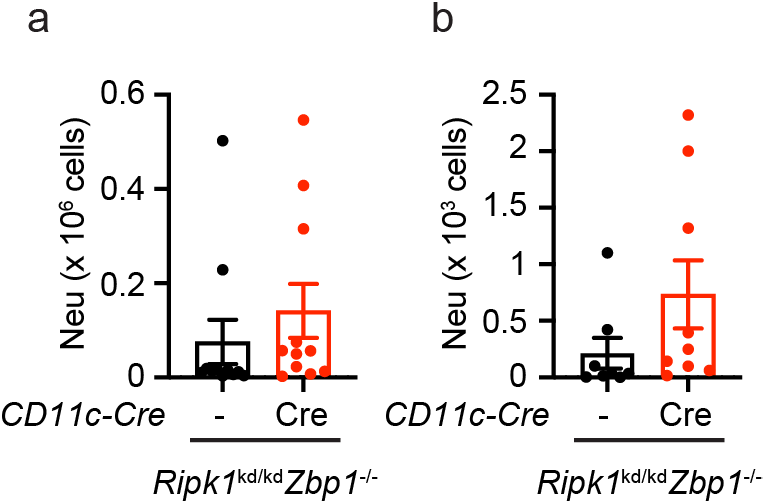
The increased neutrophil infiltration in the colon was not restored by deletion of ZBP1. Splenocytes (**a**) and colonic lamina propria cells (**b**) isolated from mice with indicated genotypes were analyzed by flow cytometry. The number of CD3^−^CD19^−^CD11c^−^CD11b^+^Ly6G^+^ neutrophils (Neu) are shown. Results are mean ± SEM (n = 8 (**a**) and 11 (**b**) for *Ripk1*^kd/kd^*Zbp1*^−/−^ mice and n = 9 (**a**) and 11 (**b**) for CD11c;*Ripk1*^kd/kd^*Zbp1*^−/−^ mice). Mice from intercross between *Ripk1*^kd/kd^*Zbp1*^−/−^ and CD11c;*Ripk1*^kd/kd^*Zbp1*^−/−^ mice were used.

### DC-specific deletion of RIPK1 confers protection against DSS-induced colitis

Patients with loss of function mutation of *RIPK1* exhibited inflammatory bowel diseases^17–19^. Although CD11c;*Ripk1*^kd/kd^ mice developed spontaneous colonic inflammation, severe colonic tissue damage was not observed. We hypothesized that stimulation with colitogenic agent might cause more severe colitis in CD11c;*Ripk1*^kd/kd^ mice than *Ripk1*^kd/kd^ mice. To test this hypothesis, we challenged the mice with dextran sulfate sodium (DSS), which induces acute damage of colonic epithelial cells and colitis-like inflammation. The mice were given 1.5% DSS water for seven days followed by regular water for seven days. During the course of experiment, body weight was measured every day to monitor severity of colitis. *Ripk1*^+/+^ mice gradually lost body weight during the first 11 days but started to recover afterwards (Fig. 5a). Loss of body weight in *Ripk1*^kd/kd^ mice was similar to that in *Ripk1*^+/+^ mice (Fig. 5a). Colon length at day 15 was similar between *Ripk1*^kd/kd^ and *Ripk1*^+/+^ mice (Fig. 5b), indicating that RIPK1 kinase activity does not affect sensitivity to DSS-induced damage. Contrary to expectation, CD11c;*Ripk1*^kd/kd^ mice were highly resistant to DSS-induced colitis and did not exhibit any body weight loss (Fig. 5c). Moreover, colon length was significantly longer in CD11c;*Ripk1*^kd/kd^ mice than *Ripk1*^kd/kd^ mice (Fig. 5d). Histological analysis revealed destructed villi and thickened submucosal layer in the colon of *Ripk1*^kd/kd^ mice but not in CD11c;*Ripk1*^kd/kd^ mice (Fig. 5e). Expression of inflammatory cytokines such as *Il6* and *Il1b*, but not *Tnf*, was significantly lower in CD11c;*Ripk1*^kd/kd^ mice than *Ripk1*^kd/kd^ mice at day 7 (Fig. 5f). In addition, the protection against DSS-induced colitis was also observed in CD11c;*Ripk1*^fl/fl^ mice (Fig. 5g). Since *Ripk1*^fl/fl^ mice were more sensitive to DSS-induced colitis, several mice had to be euthanized before day 15. Nonetheless, CD11c;*Ripk1*^fl/fl^ littermates were significantly more resistant to DSS-induced colitis than *Ripk1*^fl/fl^ mice, indicating that expression of kinase inactive RIPK1 in non-DC cell types did not contribute to the resistance to DSS. These results indicate that the scaffold-dependent but kinase-independent function of RIPK1 in DCs promotes DSS-induced colitis.

**Fig.5.**
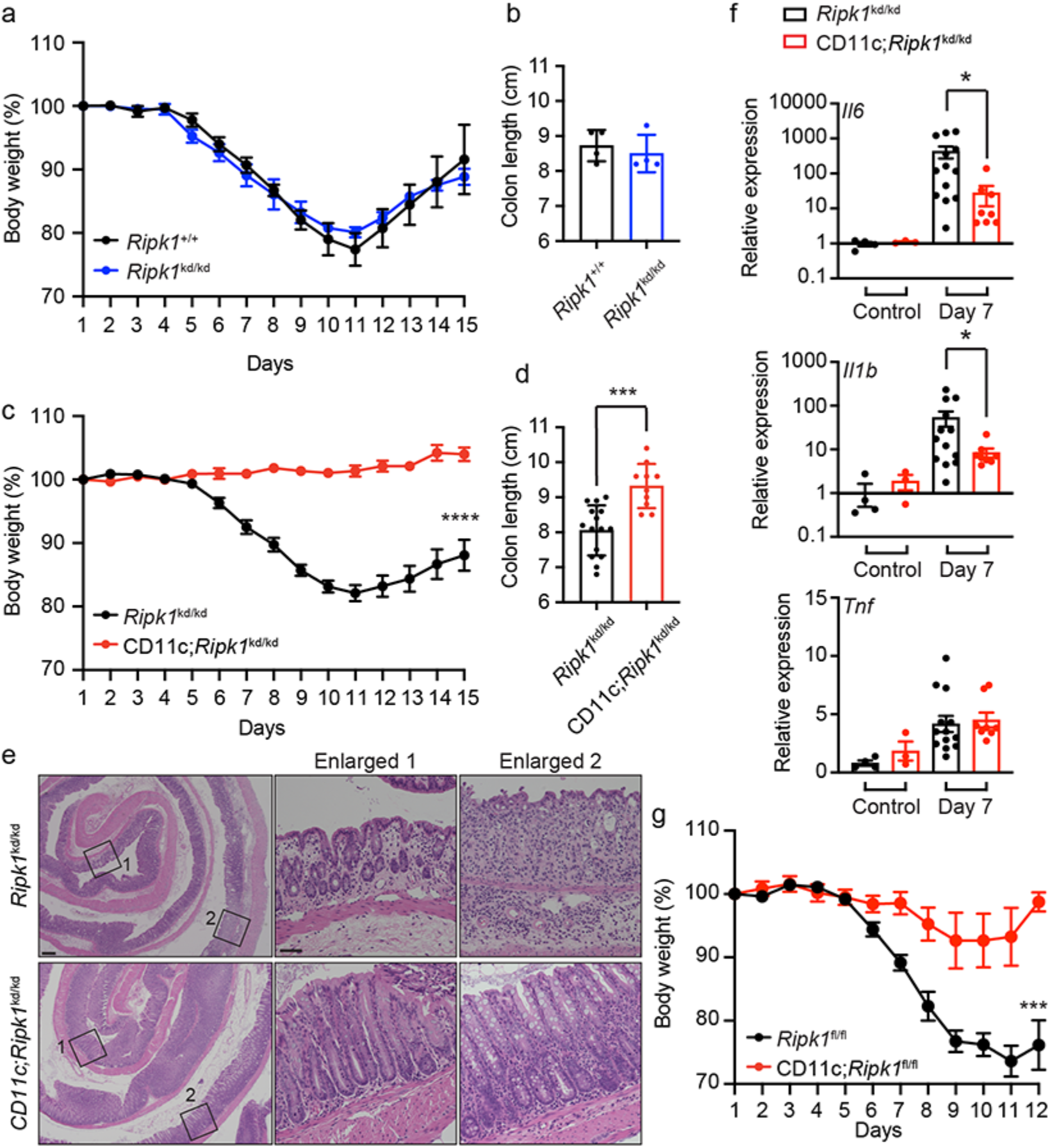
DC-specific deletion of RIPK1 confers protection against DSS-induced colitis. **a-d** Mice with indicated genotypes were treated with DSS for seven days, followed by regular water for an additional seven days. The weight at the beginning of the experiments was normalized as 100%. Daily body weight change (**a, c**) and colon length (**b, d**) on day 15 are shown. Results are mean ± SEM (n = 4 per each genotype (**a, b**), n = 15 for *Ripk1*^kd/kd^ mice and n = 10 for CD11c;*Ripk1*^kd/kd^ mice (**c, d**)). **e** H& staining of the colon from DSS-treated mice with indicated genotypes (day 7) (n = 5 per each genotype). The areas indicated by the black squares on the left (scale bars = 200 μm) were enlarged and shown in the panels on the right (scale bars = 50 μm). **f** The expression of *Il6*, *Il1b*, and *Tnf* in the colon was determined by real-time qPCR. Results are mean ± SEM (n = 3-13 per each genotype). **g** Body weight of DSS-treated mice with indicated genotypes is shown. Results are mean ± SEM (n = 10 for *Ripk1*^fl/fl^ mice and n = 7 for CD11c;*Ripk1*^fl/fl^ mice). Mice from intercross between *Ripk1*^fl/fl^ and CD11c;*Ripk1*^fl/fl^ mice were used in **g**. Mice from intercross between *Ripk1*^kd/kd^ and CD11c;*Ripk1*^kd/kd^ mice were used in others. \**p* < 0.05, \*\*\**p* < 0.001, \*\*\*\**p* < 0.0001 (two-way ANOVA in **c, g** and unpaired t test with Welch’s correction in others).

### The protective effect of DC-specific RIPK1 deletion on DSS-induced colitis was mediated by FADD

We examined whether apoptosis and/or necroptosis was involved in the protective effect of DC-specific deletion of RIPK1 on DSS-induced colitis. Similar to CD11c;*Ripk1*^kd/kd^ mice (Fig. 5a), CD11c;*Ripk1*^kd/kd^;*Ripk3*^ΔR/+^;*Fadd*^+/−^mice were highly resistant to DSS-induced colitis (Fig. 6a-c). CD11c;*Ripk1*^kd/kd^;*Ripk3*^ΔR/ΔR^;*Fadd*^+/−^ mice were also significantly protected from DSS-induced colitis compared with *Ripk1*^kd/kd^;*Ripk3*^ΔR/ ΔR;^*Fadd*^+/−^mice (Fig. 6a-c). We further found that body weight loss of CD11c;*Ripk1*^kd/kd^;*Mlkl*^−/−^ mice was significantly reduced compared to *Ripk1*^kd/kd^;*Mlkl*^−/−^ mice (Fig. 6d). These results indicate that necroptosis does not contribute to the protective effect of DC-specific RIPK1 deletion on DSS-induced colitis. In contrast, CD11c;*Ripk1*^kd/kd^;*Ripk3*^ΔR/ΔR^;*Fadd*^−/−^ and *Ripk1*^kd/kd^;*Ripk3*^ΔR/ΔR^;*Fadd*^−/−^ mice exhibited similar body weight loss and colonic inflammation in response to DSS (Fig. 6a-c). In fact, the weight loss of both *Ripk1*^kd/kd^;*Ripk3*^ΔR/ΔR^;*Fadd*^−/−^ and CD11c;*Ripk1*^kd/kd^;*Ripk3*^ΔR/ΔR^;*Fadd*^−/−^ mice were severe enough that we needed to euthanize some of the mice before day 15 (Fig. 6a). These results indicate that FADD mediates protection against DSS-induced colitis in CD11c;*Ripk1*^kd/kd^ mice. Deletion of *Zbp1* partially restored sensitivity to DSS in CD11c;*Ripk1*^kd/kd^;*Zbp1*^−/−^ mice, although these mice were still significantly protected from DSS when compared to *Ripk1*^kd/kd^;*Zbp1*^−/−^ mice (Fig. 6e), suggesting that ZBP1 might have a minor contribution to the protective effect of DC-specific RIPK1 deletion on DSS-induced colitis. Collectively, these results indicate that DC-specific RIPK1 deletion confers the protection against DSS-induced colitis through FADD-dependent apoptosis but not RIPK3-MLKL-dependent necroptosis.

**Fig.6.**
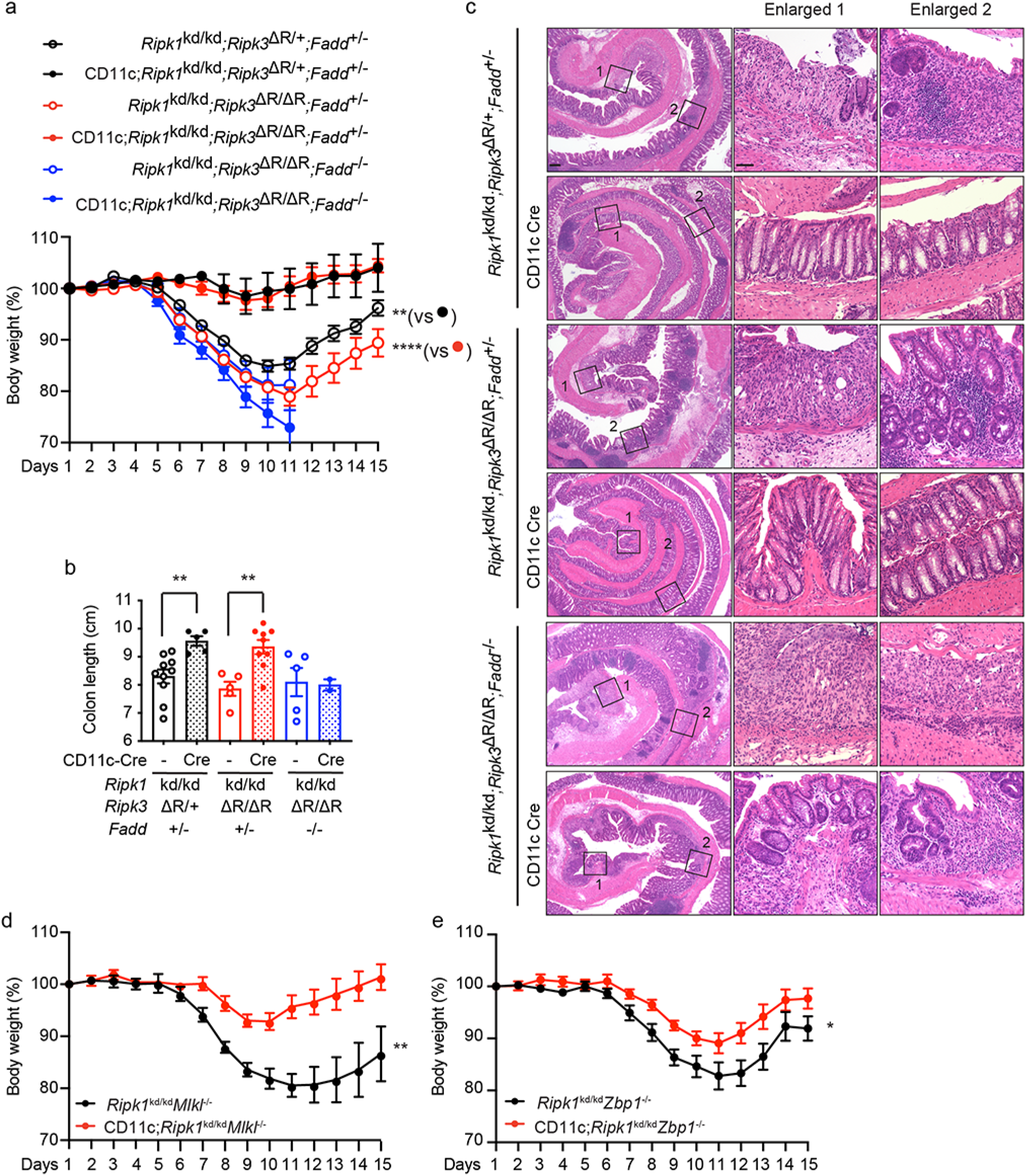
The protective effect of DC-specific RIPK1 deletion in DSS-induced colitis was abrogated by deletion of RIPK3 RHIM and FADD. **a-b** Mice with indicated genotypes were treated with DSS for seven days, followed by regular water for an additional seven days. The weight at the beginning of the experiments was normalized as 100%. Daily body weight (**a**) and colon length (**b**) at day 15 are shown. Results are mean ± SEM (n = 10 (**a, b**) for *Ripk1*^kd/kd^*Ripk3*^ΔR/+^*Fadd*^+/−^ mice, n = 5 (**a, b**) for CD11c;*Ripk1*^kd/kd^*Ripk3*^ΔR/+^*Fadd*^+/−^ mice, n = 5 (**a, b**) for *Ripk1*^kd/kd^*Ripk3*^ΔR/ΔR^*Fadd*^+/−^ mice, n = 9 (**a, b**) for *Ripk1*^kd/kd^*Ripk3*^ΔR/ΔR^*Fadd*^+/−^ mice, n = 6 (**a**) and n = 5 (**b**) for CD11c;*Ripk1*^kd/kd^*Ripk3*^ΔR/ΔR^*Fadd*^−/−^ mice, n = 6 (**a**) and 2 (**b**) for CD11c;*Ripk1*^kd/kd^*Ripk3*^ΔR/ΔR^*Fadd*^−/−^ mice). **c** H& staining of the colon from DSS-treated mice with indicated genotypes (day 15) (n = 2 per each genotype). The areas indicated by the black squares are enlarged. Scale bars: 200 μm (left panel) and 50 μm (enlarged pictures). **d, e** Body weight of DSS-treated mice with indicated genotypes is shown. Results are mean ± SEM (n = 7 for *Ripk1*^kd/kd^*Mlkl*^−/−^, n = 9 for CD11c;*Ripk1*^kd/kd^*Mlkl*^−/−^, n = 11 for *Ripk1*^kd/kd^*Zbp1*^−/−^, and n = 13 for CD11c;*Ripk1*^kd/kd^*Zbp1*^−/−^ mice). Mice from intercross between *Ripk1*^kd/kd^*Ripk3*^ΔR/ΔR^*Fadd*^−/−^ and CD11c;*Ripk1*^kd/kd^*Ripk3*^ΔR/+^*Fadd*^+/−^ mice were used in **a-c**. Mice from intercross between *Ripk1*^kd/kd^*Mlkl*^−/−^ and CD11c;*Ripk1*^kd/kd^*Mlkl*^−/−^ mice were used in **d**. Mice from the intercross between *Ripk1*^kd/kd^*Zbp1*^−/−^ and CD11c;*Ripk1*^kd/kd^*Zbp1*^−/−^ mice were used in **e**. \**p* < 0.05, \*\**p* < 0.01, \*\*\*\**p* < 0.0001 (two-way ANOVA in **a, d, e** and unpaired t test with Welch’s correction in **b**).

## Discussion

RIPK1 functions as a critical rheostat that controls cell survival and death signals. Loss of RIPK1 in mice causes early postnatal lethality due to massive cell death in multiple peripheral tissues and bone marrow failure^10, 11^. In addition, intestine-specific deletion of *Ripk1* led to lethal intestinal inflammation in mice^20, 21^. On the other hand, patients with loss of function mutation including null mutation of *RIPK1* commonly exhibits immunodeficiency and inflammatory bowel disease with shortened life expectancy ^17–19^. Although deletion of *Ripk1* in mouse epidermis caused severe skin inflammation^20^, skin disorder is not common among the *RIPK1* mutant patients. Intriguingly, hematopoietic stem cell transplantation resolved both immunodeficiency and inflammatory bowel disease of these patients ^17^. Therefore, RIPK1 in hematopoietic cells plays a key role in the maintenance of immune functions and peripheral tissue homeostasis.

In this study, we showed that DC-specific deletion of RIPK1 caused spontaneous colonic inflammation characterized by increased neutrophil infiltration. CD11c;*Ripk1*^kd/kd^ mice, which lack RIPK1 in DCs and RIPK1 kinase activity in other cell types, exhibited spontaneous colonic inflammation. The fact that *Ripk1*^kd/kd^ mice did not exhibit colonic inflammation indicates that RIPK1 kinase activity was not involved. Previously, CD11c;*Ripk1*^fl/fl^ mice were reported to develop splenomegaly and splenic inflammation, both of which are restored by deletion of either RIPK3 or MLKL^24^. This splenic phenotype was also observed in CD11c;*Ripk1*^kd/kd^ mice, indicating that loss of RIPK1 kinase activity in other cell types do not rescue the inflammation induced by RIPK1 deletion in DCs. In addition to these phenotypes, we detected increased neutrophil infiltration in the colon of CD11c;*Ripk1*^kd/kd^ mice. In contrast to the spleen, deletion of *Ripk3*, *Mlkl* or *Zbp1* did not reverse this colonic inflammation. However, co-deletion of FADD and RIPK3 RHIM in CD11c;*Ripk1*^kd/kd^ mice eliminated the colonic inflammation. These results suggest that apoptosis of colonic DCs, especially CD103^+^ DCs, might have stimulated the neutrophil infiltration through upregulation of the chemokines CXCL1 and CXCL2. Thus, our results revealed that the loss of RIPK1 could promote apoptosis or necroptosis in a tissue-specific manner. For example, RIPK1 deletion in the intestine and skin primarily caused apoptosis and necroptosis, respectively^20^. Specifically, colonic DCs are prone to die by apoptosis in the absence of RIPK1.

*Batf3*^−/−^, *Flk2*^−/−^, or *Flt3l*^−/−^ mice, all of which exhibited impaired DC development, have increased neutrophils in peripheral tissues such as liver, spleen, and lung^36^. In addition, DC depletion by administration of diphtheria toxin (DT) in mice expressing DT receptor under the regulation of CD11c promoter (CD11c-DTR mice) also caused neutrophilia^37^. Therefore, DCs are a critical cell type regulating peripheral neutrophil homeostasis. It is noteworthy that neutrophil number was increased in the bone marrow of CD11c;*Ripk1*^kd/kd^ mice while it was reduced in DT-treated CD11c-DTR mice^37^. It is likely that constitutive reduction of DC population stimulates neutrophil development in the bone marrow.

CD11c;*Ripk1*^kd/kd^ mice did not develop gross intestinal tissue destruction and diarrhea, which are characteristics of the patients with *RIPK1* deficiency. Previous studies demonstrated that mice transplanted with of *Ripk1*^−/−^ fetal liver cells and hematopoietic cell-specific *Ripk1* knockout mice also did not exhibited intestinal tissue destruction^11, 22^. These results suggest that additional factors are required to induce inflammatory bowel disease in mice. CD11c;*Ripk1*^kd/kd^ mice were unexpectedly refractory to DSS-induced colitis. This resistance was abrogated when the function of both RIPK3 and FADD, but not RIPK3, MLKL or ZBP1 alone, was disrupted. The resistance to DSS-induced colitis in the different strains of mice tested in this study correlated with an increase in colonic neutrophils. It is therefore tempting to speculate that the increased neutrophils facilitate clearance of commensal bacteria when the colonic epithelium is breached, thereby reducing inflammation triggered by commensal bacteria. In support of this model, a previous study reported that clearance of bacterial infection was enhanced when neutrophils were increased upon depletion of DCs ^37^.

We previously reported that RIPK3 promotes injury-induced cytokine production in DCs in a RHIM-dependent manner. In contrast to *Ripk1* deletion, loss of RIPK3 activity in DCs does not lead to any signs of spontaneous inflammation and that the deletion of RIPK3 RHIM in DCs exacerbated DSS-induced colitis^26–28^. Thus, although RIPK1 and RIPK3 function in synergy to promote necroptosis and detrimental inflammation, they exhibit distinct functions in colonic DCs to regulate intestinal homeostasis. Our study illustrates the multi-functional nature of RIPK1 in controlling life and death.

## Methods

### Mice

RIPK1 kinase dead mutant knock-in mice (*Ripk1*^kd/kd^) in which exon 4 was flanked by loxP sequences were obtained from GlaxoSmithKline^29^. *Ripk3*^ΔR/ΔR^ mice were generated as reported previously^28^. *Fadd*^−/−^ mice were a kind gift from J. Zhang (Thomas Jefferson University). *Mlkl*^−/−^ mice were a kind gift from J. Murphy (Walter Eliza Hall Institute). *Zbp1*^−/−^ mice were obtained from Laboratory of Animal Models for Human Diseases (National Institutes of Biomedical Innovation, Health and Nutrition). CD11c-Cre (*Itgax-cre*) transgenic mice were obtained from Jackson laboratory. *Ripk1*^fl/fl^ mice were a kind gift from M. Pasparakis (University of Cologne). All animals were housed and maintained under specific pathogen-free conditions in the animal facility at Osaka University Graduate School of Medicine, Toho University School of Medicine, and Duke University. All animal experiments were approved by the institutional animal care and use committee.

### Dextran sulfate sodium-induced colitis

DSS (MP Biomedicals, molecular weight 36,000-50,000 Da) was added to sterilized water at a concentration of 1.5% (wt/vol) and administered to age-matched mice (9-12 weeks). After the treatment with DSS for seven days, DSS water was replaced with sterilized water and the mice were sacrificed for analyses at the indicated time point. Body weight was monitored throughout the studies.

### Histological analysis

The colon was harvested and fixed with 10% formaldehyde, pH 7.0. The tissues were dehydrated and embedded in paraffin. HE staining was performed according to a standard histological procedure. For immunohistochemical staining, fixed tissues were incubated with 20% sucrose in 0.1M PB, pH 7.2 and subsequently frozen in O.C.T. Compound. The frozen tissue sections were treated with blocking solution (5% normal donkey serum in PBS) and then stained with FITC-conjugated anti-Ly6G antibody (BioLegend, 1A8). Nuclei were stained by DAPI. Finally, the tissue slides were mounted with Mowiol mounting medium. Either a BZ-X710 microscope operated by BZ-X Viewer (Keyence) or an FV1000D confocal microscope operated by FluoView software (Olympus) was used to acquire images.

### Flow cytometer

The colon was harvested and incubated in Hanks’ balanced salt solution (HBSS) containing 1 mM DTT for 10 minutes at RT. After incubating in HBSS containing 30 mM EDTA and 10 mM HEPES for 10 minutes at 37 °C twice, the tissues were minced and then incubated in digestion medium (1 mg/ml collagenase IV (Sigma, C5138), 150 μg/ml DNase II (Sigma, DN25), and 0.1 U/ml dispase (STEMCELL Technologies, #07913) in DMEM supplemented with 10% FCS) for 45 minutes at 37 °C. After vigorous shaking followed by filtration, cells were subjected to Percoll gradient centrifugation. For splenocytes, the spleen was incubated in collagenase solution (2 mg/ml Collagenase IV, 10 mM HEPES pH 7.4, 150 mM NaCl, 5 mM KCl, 1 mM MgCl_2_, 1.8 mM CaCl_2_) for 30 minutes at 37 °C. Splenocytes were subjected to erythrocyte lysis (ACK lysis buffer, 150 mM NH_4_Cl, 10 mM KHCO_3_, 100 μM EDTA). Isolated cells were incubated with anti-CD16/32 antibody (2.4G2) and subsequently stained with following antibodies: FITC-conjugated MHC class II I-Ab (eBioscience, AF6-120.1), PerCP Cy5.5-conjugated CD11b (BioLegend, M1/70), APC-conjugated CD11c (BD Biosciences, HL3), biotin-conjugated CD103 (BioLegend, 2E7), Pacific blue-conjugated CD45.2 (BioLegend, 104), APC Cy7-conjugated CD8α (TONBO Biosciences, 53-6.7), FITC-conjugated Ly6G (BioLegend, 1A8), PE Cy7-conjugated CD45.2 (BioLegend, 104), APC Cy7-conjugated CD3ε (BioLegend, 145-2C11), APC Cy7-conjugated B220 (BioLegend, RA3-6B2), PE-conjugated CD3ε (BioLegend, 145-2C11), PE Cy7-conjugated CD19 (eBioscience, eBio1D3), biotin-conjugated B220 (BD Biosciences, RA3-6B2), and APC eFluor780-conjugated streptavidin (BioLegend, 47-4317-82). For intracellular staining of RIPK1, cells stained for surface markers were fixed with 3% paraformaldehyde in PBS for 15 minutes and then permeabilized with 0.05% triton X-100 and 5% normal donkey serum in PBS for 10 minutes. Cells were incubated with anti-RIPK1 antibody (CST, 3493) or control rabbit IgG (Wako, 148-09551) for an hour followed by Alexa Fluor 594-conjugated donkey anti-rabbit IgG (H+L) (Invitrogen) for an hour. Cells were analyzed by either FACS CantoII or LSR Fortessa flow cytometers.

### Quantitative PCR

Total RNA was extracted from tissues using NucleoSpin RNA (MACHEREY-NAGEL). cDNA was synthesized using either PrimeScript reverse transcriptase (TAKARA Bio) or ReverTra Ace (TOYOBO). Real-time quantitative PCR was performed with either FastStart Universal Prove Master Mix (Roche Diagnostics) using ViiA 7 Real-Time PCR system (Applied Biosystems), FastStart SYBR Green Master (Roche Diagnostics) using LightCycler 96 (Roche Diagnostics), or Fast SYBR Green Master Mix (Applied Biosystems) using Quant Studio 3 Real-Time PCR system (Applied Biosystems). Glucose-6-phosphate dehydrogenase X-linked (G6PDX) and hypoxanthine-guanine phosphoribosyltransferase 1 (HPRT1) were used as an internal control. The following primers were used: 5’-CCACACTCAAGAATGGTCGC-3’ and 5’-TCTCCGTTACTTGGGGACAC-3’ for *Cxcl1*, F: 5’-AAAATCATCCAAAAGATACTGAACAA-3’ and R: 5’-CTTTGGTTCTTCCGTTGAGG-3’ with universal probe #66 for *Cxcl2*, 5’-CACAGAGGATACCACTCCCAA-3’and 5’-TCCACGATTTCCCAGAGAACA-3’ for *Il6*, 5’-GAGCTGAAAGCTCTCCACCTCA-3’and 5’-TCGTTGCTTGGTTCTCCTTGTAC-3’for *Il1b*, and 5’-GGTGCCTATGTCTCAGCCTCTT-3’and 5’-CGATCACCCCGAAGTTCAGTA-3’for *Tnf*.

### Statistics

Statistical analysis was performed by unpaired t test with Welch’s correction, one-way ANOVA, or two-way ANOVA using Prism 8 software. P value lower than 0.05 was considered statistically significant.

## Acknowledgments

We thank P. Gough and J. Bertin (GlaxoSmithKline) for *Ripk1*^kd/kd^ mice, J. Zhang (Thomas Jefferson University) for *Fadd*^−/−^ mice, J. Murphy (Walter Eliza Hall Institute) for *Mlkl*^−/−^ mice, M. Pasparakis (University of Cologne) for *Ripk1*^fl/fl^ mice. This study was supported by NIH grants AI119030 and AI148302 (F.K.-M.C.), the Japan Society for the Promotion of Science KAKENHI grants (K.M.: 16H06945 and 19K07399), the Takeda Science Foundation (K.M.), and GSK Japan Research Grant 2020 (K.M.).

## Author Contributions

K.M. conceived the project. K.M. and F.K.-M.C. designed experiments. K.M., C.P., K. Koyama, and S.B. performed experiments. K. Kita, R.Y., and S.K-S. assisted in maintaining the mice. K.M., C.P., H.N., A.H., Y.K., E.M., and F.K.-M.C. analyzed the data. K.M. and F.K.-M.C. wrote the manuscript. K.M. and F.K.-M.C. supervised the projects.

## Disclosure

The authors declare that they have no competing interests.

## Notes

The authors declare that they have no conflicts of interest with the contents of this article.

### Competing Interest Statement

The authors have declared no competing interest.

